# polymapR – linkage analysis and genetic map construction from F1 populations of outcrossing polyploids

**DOI:** 10.1101/228817

**Authors:** Peter M. Bourke, Geert van Geest, Roeland E. Voorrips, Johannes Jansen, Twan Kranenburg, Arwa Shahin, Richard G. F. Visser, Paul Arens, Marinus J. M. Smulders, Chris Maliepaard

**Affiliations:** Plant Breeding, Wageningen University & Research, Droevendaalsesteeg 1, 6708 PB Wageningen, The Netherlands.; Horticulture and Product Physiology, Wageningen University & Research, Droevendaalsesteeg 1, 6708 PB Wageningen, The Netherlands; Biometris, Wageningen University & Research, Droevendaalsesteeg 1, 6708 PB Wageningen, The Netherlands.; Van Zanten Breeding B. V., Lavendelweg 15, 1435 EW, Rijsenhout, The Netherlands.

**Keywords:** autopolyploid genetic map, triploid, tetraploid, hexaploid, partial preferential chromosomal pairing, segmental allopolyploid

## Abstract

**Motivation:** Polyploid species carry more than two copies of each chromosome, a condition found in many of the world’s most important crops. Genetic mapping in polyploids is more complex than in diploid species, resulting in a lack of available software tools. These are needed if we are to realise all the opportunities offered by modern genotyping platforms for genetic research and breeding in polyploid crops.

**Results:** polymapR is an R package for genetic linkage analysis and integrated genetic map construction from bi-parental populations of outcrossing autopolyploids. It can currently analyse triploid, tetraploid and hexaploid marker datasets and is applicable to various crops including potato, leek, alfalfa, blueberry, chrysanthemum, sweet potato or kiwifruit. It can detect, estimate and correct for preferential chromosome pairing, and has been tested on high-density marker datasets from potato, rose and chrysanthemum, generating high-density integrated linkage maps in all of these crops.

**Availability and Implementation:** polymapR is freely available under the general public license from the Comprehensive R Archive Network (CRAN) at http://cran.r-project.org/packages=polymapR.

**Contact:** Chris Maliepaard or Roeland E. Voorrips

## Introduction

In recent years there has been an acceleration of progress in the understanding of the genetics underlying important traits in autopolyploid species. This has been to a large extent due to developments in high-density genotyping platforms for single nucleotide polymorphism (SNP) markers, which have found increasing application in polyploids. For example, high-density SNP arrays have been developed in potato (Felcher et al., 2012; Vos et al., 2015), rose (Koning-Boucoiran et al., 2015), alfalfa (Li et al., 2014) and chrysanthemum (van Geest et al., 2017a), enhancing the scope for genetic studies in these species.

In polyploid species, as opposed to diploids, co-dominantly scored markers can possess multiple classes in the heterozygous condition, usually termed marker “dosage”. In a tetraploid there are five possible dosage classes of a bi-allelic SNP marker, namely nulliplex with a dosage 0 for one of the alleles, simplex with dosage 1, duplex with dosage 2, triplex with dosage 3, and quadruplex with dosage 4. In a hexaploid, the number of dosage classes at a bi-allelic locus rises to seven. Various software have been developed to convert the signal from *e.g.* SNP arrays into these discrete dosage calls for polyploids, such as fitTetra (Voorrips et al., 2011) or ClusterCall (Schmitz Carley et al., 2017).

Genetic linkage maps have traditionally been used for both exploratory trait mapping (often termed QTL analysis) and the subsequent fine mapping of traits, as well as for assisting genome assembly efforts by guiding the integration and orientation of contigs. High-density linkage maps may also improve our understanding of the chromosomal composition and genetics of polyploid species, uncovering such phenomena as double reduction or partially-preferential chromosome pairing. In many polyploid species which lack reference genome sequences, linkage maps are also a (vital) first genomic description of that species.

Despite the importance of both linkage maps and polyploid species, there are still relatively few software tools available for polyploid linkage map construction. Allopolyploid species showing disomic inheritance can be treated (genetically-speaking) as diploids, with a wide range of software options available. In the case of polysomic polyploids (autopolyploids and segmental allopolyploids), the options available to the research community are limited. Probably the most well-known autopolyploid mapping software is TetraploidMap (Hackett and Luo, 2003; Hackett et al., 2007), which has been used in studies of various autotetraploid species such as potato, alfalfa, rose and blueberry (e.g. (Bradshaw et al., 2008; Robins et al., 2008; Gar et al., 2011; McCallum et al., 2016)). Recently, its successor TetraploidSNPMap (TSNPM) has been released to accommodate high-density marker data from SNP arrays (Hackett et al., 2017). However, it can only handle autotetraploid datasets and provides a graphical user interface for the Windows platform only. Linkage studies in species exhibiting strong preferential chromosomal pairing or other ploidy levels are not currently possible using this software. An alternative polyploid mapping software is the PERGOLA package in R (Grandke et al., 2017). However, this software has been developed for use with F2 or backcross populations from homozygous parents only. In many cases, either due to inbreeding depression or the difficulties imposed by polysomic inheritance, F1 populations from two heterozygous parents are typically used instead.

In short, there is currently no software which can perform linkage mapping at various ploidy levels under a variety of inheritance models for outcrossing species using dosage-scored marker data. Here we present polymapR, an R package (R Core Team, 2016) for linkage mapping in outcrossing polyploid species which can generate linkage maps for polysomic triploids, tetraploids and hexaploids, accommodating either fully tetrasomic or mixed meiotic pairing behaviour (segmental allopolyploidy) at the tetraploid level. Its modularity will facilitate its adaption to other marker genotyping technologies or ploidy levels in the future.

## System and methods

The polymapR pipeline consists of four parts – data inspection, linkage analysis, linkage group assignment and marker ordering, which are detailed below. A description of the functions within polymapR is described in the vignette which accompanies the package, going through all the steps in a typical mapping project. For consistency and simplicity, all examples mentioned here describe a tetraploid cross.

### 1. Data inspection, filtering and preparation for linkage analysis

The input data for polymapR is dosage-scored marker data, available from a number of packages such as fitTetra (Voorrips et al., 2011), fitPoly *(in preparation,* Voorrips et al.) or ClusterCall (Schmitz Carley et al., 2017). Both fitTetra and ClusterCall are limited to tetraploid data whereas fitPoly can work over multiple ploidy levels. Regardless of how it is generated, the input dosage-scored marker data should consist of a column of marker dosages for the mother, one for the father followed by a column for each of the offspring of the F1 cross. Checks for marker skewness and shifted markers (when dosage scores are shifted by a fixed amount) are currently provided in polymapR from a suite of tools developed for the fitTetra package (Voorrips et al., 2011).

The next step in data preparation is the conversion of marker dosages to their simplest form, such that the sum of the parental dosage scores is minimised. There are two possible conversions – a relabelling of the reference and alternative allele in both parents, or a single-parent relabelling if the other parent is homozygous. Marker conversions are performed to reduce the number of marker segregation classes for the linkage analysis (which is directed according to the parental dosages), but have no effect on the pairwise results. In a tetraploid there are nine fundamental segregation types, rising to nineteen for a hexaploid. Identifiable double reduction scores are preserved during conversion (*e.g.* a dosage of 0 from a triplex x nulliplex (3x0) marker becomes a dosage of 2 in its converted form as a simplex x nulliplex (1x0) marker), allowing an investigation of double reduction post-mapping. Any impossible scores (like a dosage of 3 or 4 from a 1x0 marker) are made missing.

High-quality data facilitates the generation of high-quality maps. One indication of poor data quality is a high proportion of missing values. The user may choose to screen out markers or individuals with more than a desired rate of missing values (by default up to 10% is tolerated), or duplicate individuals. Identical markers, which often occur in high-density marker datasets with limited population sizes and hence a limited number of recombination events, can be identified and reduced to one representative marker for the mapping steps, and reintegrated later. A principal component analysis (PCA) can also be performed and visualised, which may highlight some unwanted structure in the population (for example due to pollination from an unknown external pollen parent or from self-pollination) or outlying individuals (for example because of admixture).

### 2. Linkage analysis

#### 2.1 Linkage analysis under a polysomic model

In autopolyploid species with polysomic inheritance, it is possible to model meiotic pairing structures as random bivalents or multivalents. In practice, both pairing structures tend to occur, with a relatively low frequency of multivalents in stable autopolyploids (Santos et al., 2003; Bomblies et al., 2016). The main consequence of multivalent formation from a genetic perspective is the phenomenon of double reduction, where two segments of a particular homologue can end up in the same gamete and become transmitted together to F1 offspring. It has been demonstrated that double reduction introduces some bias in recombination frequency estimates under a random bivalent model. This can be safely ignored if the rate of quadrivalent pairing is low (Bourke et al., 2015; Bourke et al., 2016).

Under a random bivalent model, there are three possible bivalent pairing conformations in a tetraploid. In general, for any even ploidy *p* = 2*n* there are 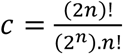 possible bivalent pairing conformations to be considered. Given any pair of marker loci with unknown recombination frequency *r*, we consider the contribution of recombinant homologues with a within-bivalent probability of 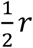 and non-recombinant homologues with a within-bivalent probability of 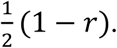 In cases where recombinants and non-recombinations cannot be distinguished, both are assigned a probability of 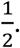 Assuming random pairing, the probability of any particular pairing configuration is 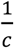 (in the case of preferential pairing, we introduce a preferential pairing factor to model deviations from randomness here).

The expected frequency of each offspring class *n_ij_* (0 ≤ *i*,*j* ≤ 2*n*) is first summed over all *c* bivalent conformations:

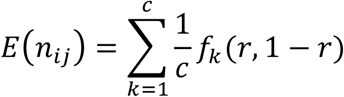

where *f_k_* (*r*, 1 − *r*) denotes a function of *r* and 1 − *r*, dependant on the marker combination considered. Given these expected frequencies, we relate them to the observed counts of individuals in each class *0*(*n_ij_*) to yield the likelihood function *𝓛*(*r*):

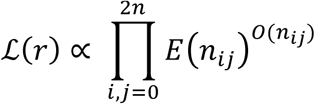

The likelihood equation results from equating the first derivative of the log of the likelihood function with zero:

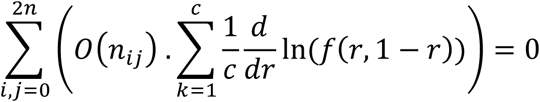

In cases where no analytical solution exists, we use Brent’s algorithm (Brent, 1973) to numerically maximise the log likelihood function in the bounded interval 0 ≤ *r* ≤ 0.5. For any pair of markers there are a number of possible phases between these markers to consider, which describe the physical linkage between marker alleles. In the case of a pair of duplex x nulliplex (2x0) markers, these phases are termed “coupling”, “mixed” and “repulsion” (Figure 1.a). As the phase between markers is initially unknown, we must compute expressions for each of the possible phases, and select the most likely as the phase for which 0 ≤ *r* ≤ 0.5, which maximises the log of the likelihood (Hackett et al., 2013).

**Figure 1.**
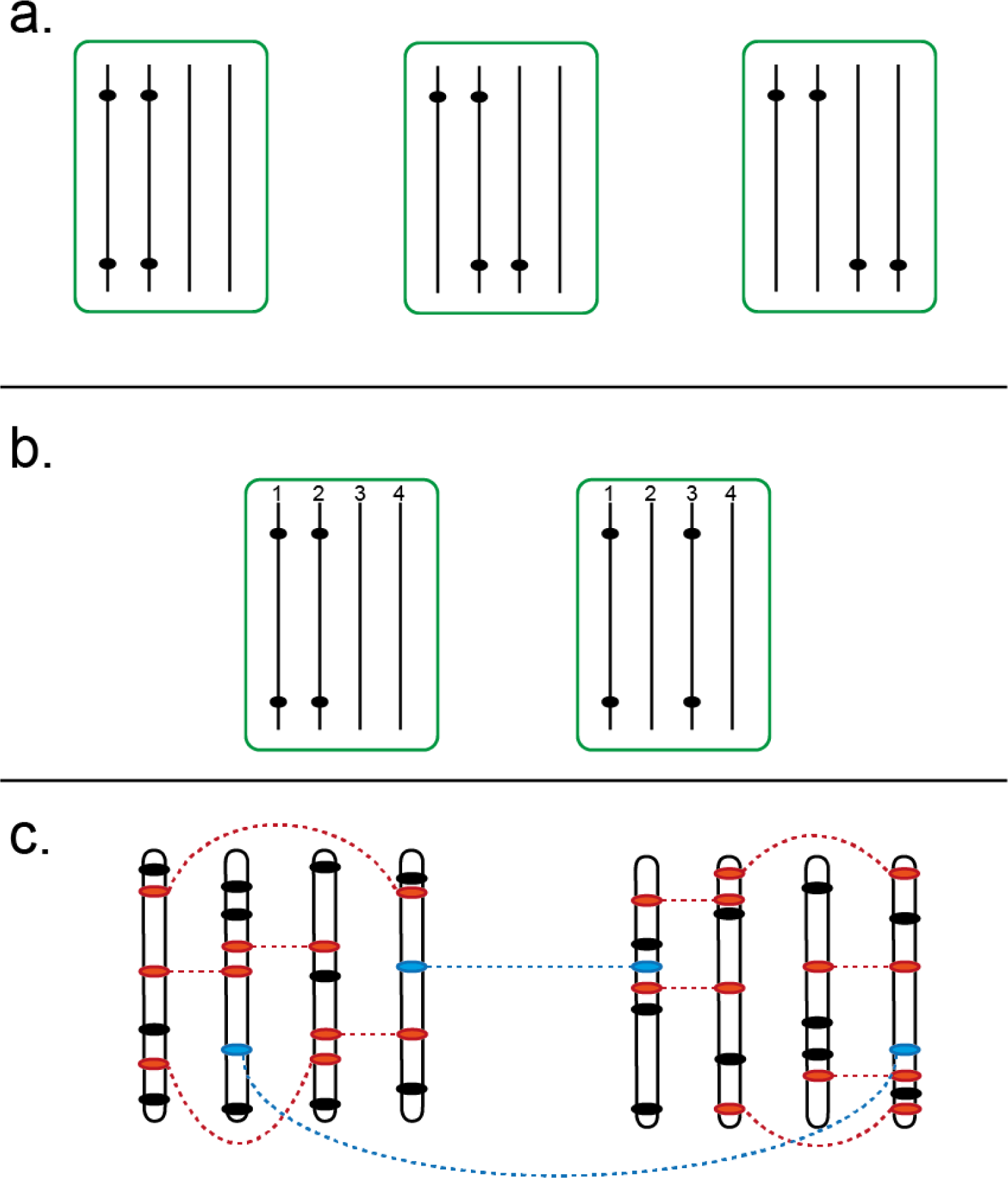
Phase considerations and clustering strategy in a tetraploid. **1.a.** The three phases considered for a pair of 2x0 markers, from left to right, “coupling”, “mixed” and “repulsion”; **1.b.** In the case of preferential pairing between homologues 1–2 and 3–4, we must consider two separate types of coupling phase, either coupling within preferential bivalents (left) or coupling between preferential bivalents (right). In the extreme case of an allotetraploid, this distinction could also be termed “subgenome-specific” versus “subgenome-straddling”. *1.c.* Simplex x nulliplex (1x0) markers (solid black dots) uniquely define homologous chromosomes and are initially clustered together. Higher-dosage marker types such as duplex x nulliplex (2x0) markers (red dots) provide linkage associations between simplex x nulliplex homologues, helping to identify chromosomal linkage groups. Cross-parental markers such as simplex x simplex (1x1, blue dots) can also link these groups together, leading to consistent linkage group numbering across parents.

Finally, we also compute the logarithm of odds (LOD) score, which provides a useful measure of the confidence in the estimate and is used for both marker clustering and marker ordering:

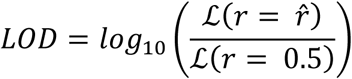

where 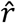 is the maximum likelihood estimate of *r*.

#### 2.2 Linkage analysis in the presence of preferential chromosomal pairing

In certain polyploid species the meiotic pairing is neither fully random nor fully partitioned into exclusively-pairing subgenomes, a situation described as segmental allopolyploidy (Stebbins, 1947). Regardless of the underlying mechanism, the result of preferential pairing is that both the segregation ratios and the co-inheritance of marker alleles are affected. In the example of a 2x0 marker introduced earlier, the expected segregation ratio in a polysomic autotetraploid is 1:4:1. With increasing preferential pairing, this ratio will approach 1:2:1 in the case of subgenome-straddling markers (Figure 1.b right), or approach non-segregation in the case of subgenome-specific markers (Figure 1.b left).

In order to model this behaviour, we introduce a preferential pairing parameter *ρ*, such that (in the case of a tetraploid) the probability of the chromosome pairing configuration 1–2 / 3–4 is 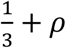 and the probability of pairing configurations 1–3 / 2–4 and 1–4 / 2–4 is 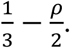 Attempting to model preferential pairing at higher ploidy levels introduces further complications; Zhu et al. (2016) have proposed a solution for hexaploids by introducing three preferential pairing parameters *θ_1_*,*θ_2_*,and *θ_3_* to model deviations in bivalent configurations 1–2, 3–4 and 5–6
respectively, with all other configurations having a probability of 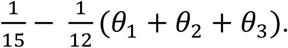 In our software, we have not yet attempted to model segmental allohexaploidy, and confine our attention to the tetraploid level for now.

We do not simultaneously estimate *ρ* and *r*, which can lead to an over-estimation of the preferential pairing parameter (Wu et al., 2002). Instead, we estimate the chromosome-wide strength of preferential pairing after map construction and thereafter correct the pairwise recombination frequency estimates to revise the maps. A robust method of preferential pairing detection and estimation is to use inheritance probability estimates such as those provided by TetraOrigin (Zheng et al., 2016); in polymapR we offer a simpler likelihood-based approach which uses closely-linked repulsion marker pairs to test for deviations from random pairing and simultaneously estimate the strength of this deviation:

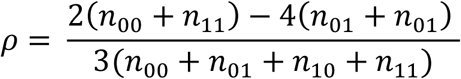

where *n*_01_ is the number of offspring with a dosage 0 at marker A and 1 at marker B *etc.*

Given a parent-and chromosome-specific estimate for the preferential pairing factor *ρ*, we modify the expression for the expected frequency of individuals in marker class *n_ij_* of a tetraploid as follows:

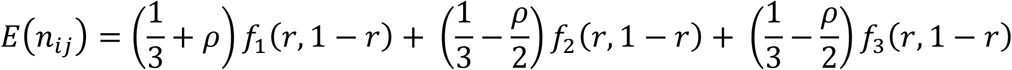

Due to the lack of symmetry, we must consider all possible conformations *within* each phase, an example of which is shown in Figure 1.b. The procedure for estimating *r* and LOD remain otherwise the same. The inclusion of preferential pairing imposes an extra computational burden as each phase can have up to four sub-phase conformations, all of which are calculated prior to selection of the most likely phase and its associated *r* and LOD score.

Finally, in both the case of random and preferential pairing, linkage calculations can be run in parallel (using the packages doParallel and doSNOW (Revolution Analytics and Weston, 2014a, b)) on any Windows or Unix-like multi-core desktop computer resulting in significant time-savings. High-density marker datasets with tens of thousands of markers can be processed in a few hours.

#### 3. Linkage group assignment

In diploid studies, the term linkage group is loosely synonymous with the term chromosome. In autopolyploids two levels of linkage group exist – homologue groups and integrated chromosomal groups. The first step in linkage group assignment is to cluster the 1x0 linkage data into homologue groups, for which we currently use the R package igraph (Csardi and Nepusz, 2006). Clustering is performed using the pairwise linkage LOD scores, although the LOD for independence can be used if desired, which may be more robust with skewed marker data (Van Ooijen and Jansen, 2013).

A number of visual aids are provided to assist in clustering (Figure 2). In general, clustering should be performed over a suitable range of LOD thresholds (*e.g.* from LOD 3 to 10) in order to inform the choice of LOD score to partition the data into both homologues and chromosomes (Figure 2.a, b). If chromosome and homologue clusters cannot be readily identified using 1x0 markers alone, coupling-phase homologue clusters are first identified at a high LOD and later re-connected into chromosomal clusters using a higher-dose marker type (Figure 1.c). Visualisations help display the strength of associations between homologues (Figure 2.c).Occasionally homologues may split apart; various possibilities to merge these fragments are provided (Figure 2. d, e, f).

**Figure 2.**
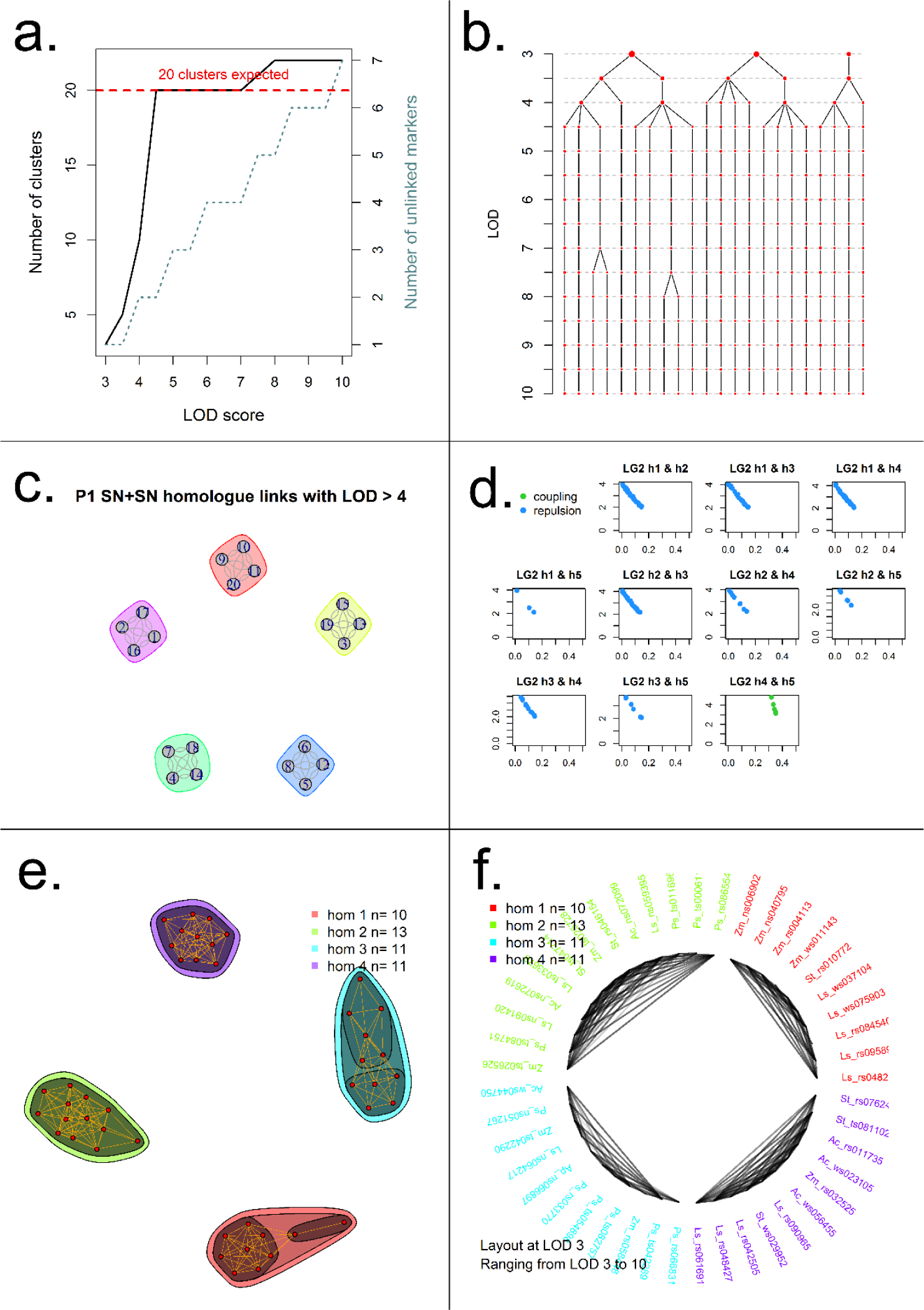
Example visualisations produced by polymapR to facilitate linkage group identification and marker clustering. **2.a.** As LOD score is increased, the number of 1x0 clusters increases, as does the number of single-marker clusters (unlinked markers). For a given ploidy and chromosome base number, the expected number of (homologue) clusters is also shown. **2.b.** Alternative representation of 2.a which shows the splitting of each cluster as the LOD score is increased. In this example, five chromosomal clusters are identified at LOD 3.5, which split into four homologue clusters between LOD 4.5 and 7. **2.c.** Using linkage to other marker segregation types such as 2x0 markers, homologue clusters can be associated into chromosomal clusters, if this was not achieved using 1x0 data alone. Here, five chromosomes are represented. **2.d.** If homologues fragment, cross-homologue phase information can help determine which fragments to merge. Here, homologues 4 and 5 show only coupling-phase linkage and should therefore be joined as a single homologue. **2 e.** Alternative approach to merge fragments showing network of linkages over a range of LOD scores. Here, four homologues were successfully identified and merged directly. **2.f.** Alternative representation of 2.e, showing these connections in a circular format instead.

In the case of triploid populations, the phasing approach differs between the diploid and tetraploid parents: for the diploid parent, phasing can be achieved directly using the phase assignment from the linkage analysis. Following the definition of the chromosome and homologue structure using the 1x0 markers, all other marker segregation classes are assigned to both homologues and chromosomes using their linkage to these markers, generating the final phase assignment of all marker types.

### 4. Marker ordering

One of the challenges of marker ordering and map construction in autopolyploid species using marker dosages is the variable accuracy of recombination frequency estimates which must be integrated somehow. Ordering algorithms which only use unweighted recombination frequency estimates are unlikely to find an optimal map order, as there is no distinction between equal estimates of *r* from situations with vastly different information contents and variances. A thorough description of this issue is provided in Preedy and Hackett (2016). Within the polymapR package, marker ordering can be achieved in two ways – either using the weighted regression algorithm as originally developed by Piet Stam (Stam, 1993) and implemented in JoinMap (Van Ooijen, 2006) and now in polymapR, or to use the multi-dimensional scaling algorithm as implemented in the MDSmap package (Preedy and Hackett, 2016). Given the computation efficiency of the MDS algorithm, in almost all circumstances this will be the preferred choice. Identical markers that were originally set aside can be added back to the final maps after marker ordering is complete.

## Implementation

### Software output - final linkage maps

The final output of the polymapR package is a phased integrated map. Maps can either be generated per homologue or per chromosome, facilitating the definition of haplotypes within a population. A record is kept in a log file of any markers that were removed at any stage during the procedure, as well as logging the function calls that generated each step, improving project reproducibility and later reporting. Visualisations are provided throughout the mapping procedure, facilitating the diagnosis of issues as well as summarising the results. An example of an integrated map with five chromosomes, generated using the sample data provided with the package, is shown in Figure 3.a. Phased linkage maps, giving the position of the SNP alleles on each parental homologue are also generated, as visualised in Figure 3.b for a triploid species. polymapR also generates input files for TetraOrigin (Zheng et al., 2016) which can calculate IBD probabilities for the population, useful for QTL analysis.

### Application of polymapR to real data

Various developmental versions of the polymapR package have been used for linkage map construction in potato, rose and chrysanthemum (Bourke et al., 2016; Vukosavljev et al., 2016; Bourke et al., 2017; van Geest et al., 2017b). The current version brings together all the capacities developed previously, while extending the algorithm to triploid populations as well (produced in a tetraploid x diploid cross). Cross-ploidy hybrids are commonly produced in ornamental breeding, as well as in certain fruit species such as watermelon (*Citrullus lanatus* var. *lanatus*) or grape (*Vitis vinifera*) to generate seedless fruit (Acquaah, 2012). polymapR is applicable to a wide range of commercially-important crop species such as potato, leek, alfalfa, blueberry, chrysanthemum, sweet potato and kiwifruit, as well as the myriad of cross-ploidy populations developed in ornamental and fruit breeding programmes.

**Figure 3.**
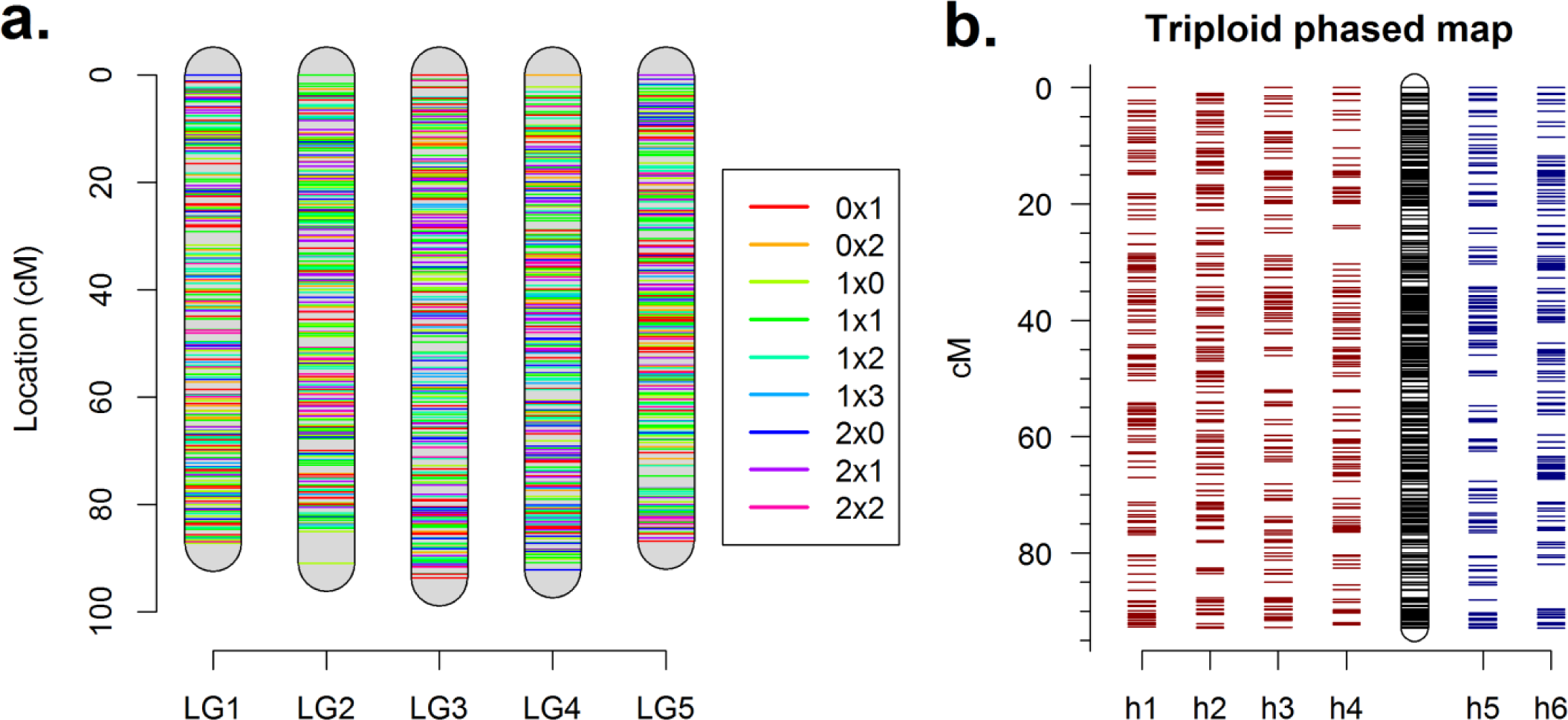
Linkage map visualisations of polymapR. **3.a.**Integrated chromosomal linkage maps generated using the sample tetraploid dataset provided with the package, with each marker segregation type coloured differently. **3.b.** Phased homologue-specific maps of a single chromosomal linkage group from a triploid dataset (simulated with PedigreeSim (Voorrips and Maliepaard, 2012)). Maternal homologue maps (h1 – h4) from the tetraploid parent are shown in red, and paternal homologue maps (h5 – h6) from the diploid parent are shown in blue, with the integrated chromosomal map in black.

## Discussion

### Comparison with other polyploid mapping software

The range of options for linkage mapping in autopolyploid species is quite limited. We compared the performance and applicability of polymapR with two alternative software, TetraploidSNPMap and PERGOLA.

### TetraploidSNPMap (TSNPM)

TSNPM possesses a graphical user interface for Windows, uses optimised routines for marker clustering and offers interactive cluster plots for linkage group assignment. It goes beyond linkage map construction to compute IBD probabilities and perform QTL interval mapping as well. Given that polymapR uses the same random bivalent pairing assumption and the same ordering algorithm (MDSmap (Preedy and Hackett, 2016)), we did not expect much difference in output. Using the sample tetraploid dataset provided with polymapR (with 3000 markers over 5 chromosomes and 207 F1 individuals, including 7 pairs of duplicate individuals), polymapR produced phased maps within 24 minutes on an Intel i7 desktop with 16 Gb RAM; TSNPM took 5 minutes, but took another 10 minutes to phase (so a total of 15 minutes were needed). However, the phased output of TSNPM is more difficult to interpret than that of polymapR and would likely require extra time for curation. The maps themselves were remarkably similar in terms of numbers of mapped markers, map length and marker order (Supplementary Figure 1).

Marker phasing in polymapR is automatic, by selecting phase based on the counts of significant linkages to 1x0 homologue clusters and ignoring any spurious linkages that go against the general trend. On the other hand, phase assignment seems to (generally) require manual intervention in the TSNPM pipeline. Despite its computational efficiency, TSNPM has also set an upper limit of 8000 SNP markers, and the maximum mapping population size is currently 300 F1 individuals. polymapR sets no limits on marker numbers or population sizes, employing parallel processing to help speed up calculations for large datasets. Duplicated markers are initially binned (also possible in TSNPM) and identical individuals are merged (this feature was missing from TSNPM) to avoid needless calculations. Overall, the main difference between TSNPM and polymapR appears to be in applicability: polymapR can analyse autotriploid, autotetraploid, autohexaploid as well as segmental allotetraploid data, whereas TSNPM is currently confined to autotetraploid data. polymapR is also cross-platform given that it is written in R (R Core Team, 2016).

### PERGOLA

The PERGOLA package in R has been developed for F2 or backcross populations from an initial cross between homozygous parents. Such a situation is highly unusual for most polysomic polyploids, since inbreeding requires many more generations before homozygosity is reached compared to a diploid or disomic polyploid. In a polysomic hexaploid for example, it would take 25 generations of selfing an F1 individual before 90% homozygosity is reached (ignoring the effects of double reduction (Haldane, 1930)). The applicability of the PERGOLA software to real populations in polysomic polyploids is therefore limited.

Despite the highly unusual type of population, we simulated a small F2 dataset of selfed F1 individuals randomly chosen from a cross between two inbred parental lines using PedigreeSim (Voorrips and Maliepaard, 2012), leading to a marker dataset of 500 duplex x duplex markers over 5 chromosomes. The calculation of recombination frequencies took a mere 3.54 seconds in PERGOLA, in comparison to 28 minutes using polymapR (on a single core; using 6 cores this step took 8 minutes). However, for polymapR this particular marker combination is complex, with nine possible phase combinations in the parents to be separately calculated per marker pair, and with extremely complicated likelihood functions for each phase (all 25 dosage combinations are possible in a tetraploid, from *n_00_* to *n_44_*). It is therefore a somewhat unfair comparison, as PERGOLA labours under no such “generalist” difficulties. Phase considerations are trivial and therefore ignored by PERGOLA because of their simplistic population assumptions. If such populations could be generated, PERGOLA would produce excellent maps. In our test, PERGOLA identified all five chromosomes, with near perfect marker order in each, although the map lengths were inflated – from 200 cM using the Kosambi mapping function to 400 cM using Haldane’s (when 100 cM was expected). polymapR also produced near-perfect maps with map-lengths of approximately 90 cM using Haldane’s mapping function. The polymapR package can handle data from both cross-pollinating *and* inbred populations whereas PERGOLA cannot, but given the performance difference, PERGOLA would appear to be the software of choice for inbred polyploid populations, should they be developed.

### Concluding remarks

The development and release of polymapR comes at a time when there is increasing need for tools to perform genetic analysis in polyploids. Understanding the genetic control of important biological traits in polyploid species will have a large impact on plant breeding (or in the case of certain salmonid fish, animal breeding as well), facilitating the adoption of genomics-driven breeding decisions such as marker-assisted selection or genomic prediction into breeding programs. For these advances to take place, high-density and accurate maps showing the relative position of markers on chromosomal groups are needed – which is precisely what polymapR delivers.

## Acknowledgements

The authors wish to thank Dr. Katherine Preedy and Dr. Christine A. Hackett (Biomathematics Scotland, Dundee, Scotland) for providing a developmental version of the MDSmap scripts before their package became publically available, and to Dr. Johan van Ooijen (Kyazma B.V. Wageningen, The Netherlands) for helpful comments. This work was supported through the TKI projects “A genetic analysis pipeline for polyploid crops” (project number B0-26.03-002-001) and “Novel genetic and genomic tools for polyploid crops” (project number BO-26.03-009-004).

## Supplementary Information

Supplementary material is available

**Supplementary Figure 1.**
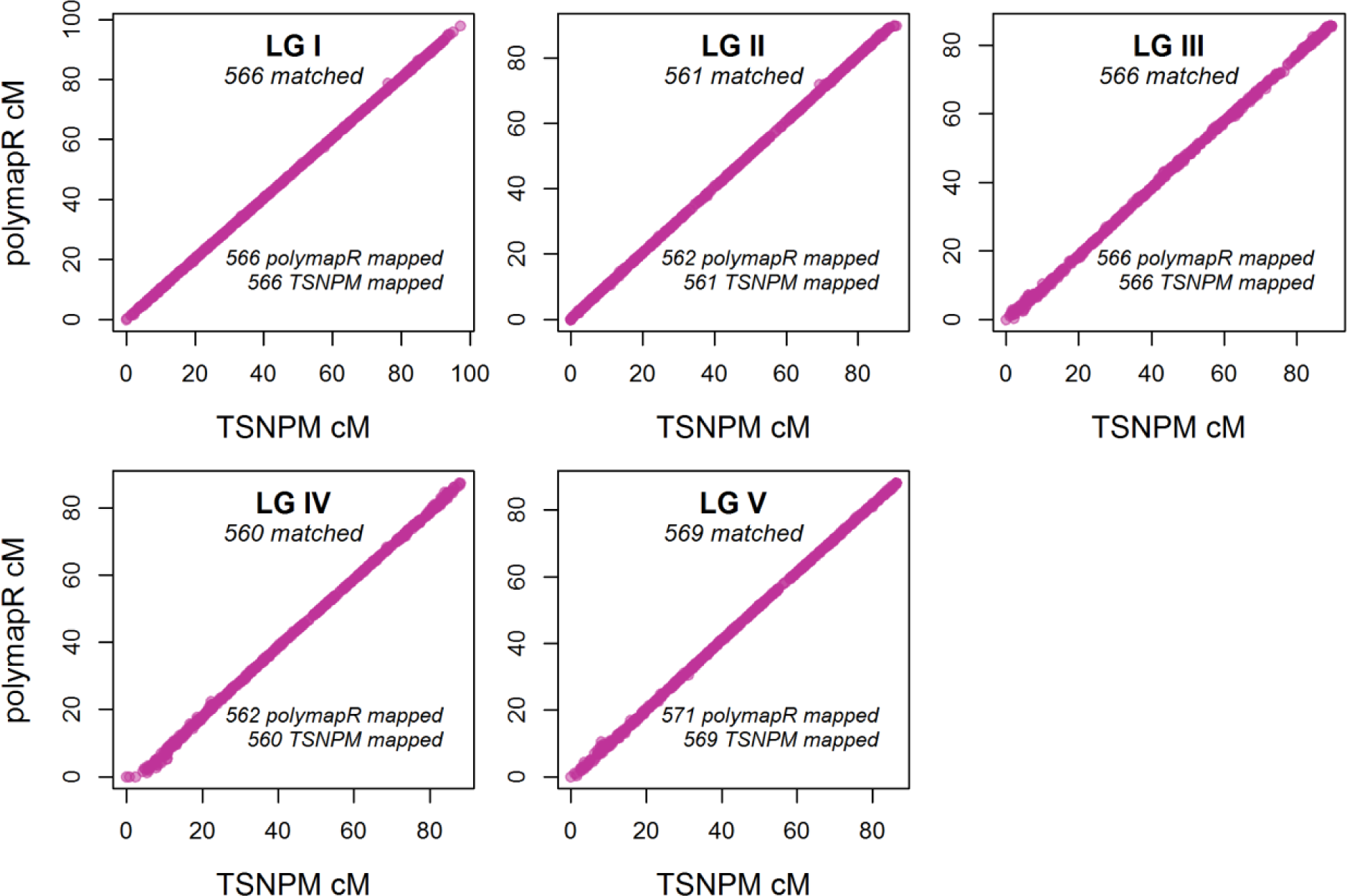
Comparison of final linkage maps produced by polymapR (y-axis) and TetraploidSNPMap (x-axis) using the sample dataset provided with the polymapR package. For all linkage groups, the marker order, map length and the number of mapped markers were almost identical between both software

## References

Acquaah, G. (2012). Principles of Plant Genetics and Breeding. (Wiley-Blackwell).

Bomblies, K., Jones, G., Franklin, C., Zickler, D., and Kleckner, N. (2016). The challenge of evolving stable polyploidy: could an increase in “crossover interference distance” play a central role? Chromosoma 125, 287–300.

Bourke, P.M., Voorrips, R.E., Visser, R.G.F., and Maliepaard, C. (2015). The Double Reduction Landscape in Tetraploid Potato as Revealed by a High-Density Linkage Map. Genetics 201, 853–863.

Bourke, P.M., Voorrips, R.E., Kranenburg, T., Jansen, J., Visser, R.G., and Maliepaard, C. (2016). Integrating haplotype-specific linkage maps in tetraploid species using SNP markers. Theoretical and Applied Genetics 129, 2211–2226.

Bourke, P.M., Arens, P., Voorrips, R.E., Esselink, G.D., Koning-Boucoiran, C.F.S., van ‘t Westende, W.P.C., Santos Leonardo, T., Wissink, P., Zheng, C., Van Geest, G., Visser, R.G.F., Krens, F.A., Smulders, M.J.M., and Maliepaard, C. (2017). Partial preferential chromosome pairing is genotype dependent in tetraploid rose. The Plant Journal 90, 330–343.

Bradshaw, J.E., Hackett, C.A., Pande, B., Waugh, R., and Bryan, G.J. (2008). QTL mapping of yield, agronomic and quality traits in tetraploid potato *(Solanum tuberosum* subsp. *tuberosum)*. Theoretical and Applied Genetics 116, 193–211.

Brent, R. (1973). Algorithms for minimizing without derivatives (Prentice-Hall, Englewood Cliffs, NJ.).

Csardi, G., and Nepusz, T. (2006). The igraph software package for complex network research. InterJournal, Complex Systems 1695, 1–9.

Felcher, K.J., Coombs, J.J., Massa, A.N., Hansey, C.N., Hamilton, J.P., Veilleux, R.E., Buell, C.R., and Douches, D.S. (2012). Integration of two diploid potato linkage maps with the potato genome sequence. PLoS ONE 7, e36347.

Gar, O., Sargent, D.J., Tsai, C.-J., Pleban, T., Shalev, G., Byrne, D.H., and Zamir, D. (2011). An autotetraploid linkage map of rose *(Rosa hybrida)* validated using the strawberry *(Fragaria vesca)* genome sequence. PLoS ONE 6, e20463.

Grandke, F., Ranganathan, S., van Bers, N., de Haan, J.R., and Metzler, D. (2017). PERGOLA: fast and deterministic linkage mapping of polyploids. BMC Bioinformatics 18, 12.

Hackett, C., and Luo, Z. (2003). TetraploidMap: construction of a linkage map in autotetraploid species. Journal of Heredity 94, 358–359.

Hackett, C.A., McLean, K., and Bryan, G.J. (2013). Linkage Analysis and QTL Mapping Using SNP Dosage Data in a Tetraploid Potato Mapping Population. PLoS ONE 8, e63939.

Hackett, C.A., Milne, I., Bradshaw, J.E., and Luo, Z. (2007). TetraploidMap for Windows: linkage map construction and QTL mapping in autotetraploid species. Journal of Heredity 98, 727–729.

Hackett, C.A., Boskamp, B., Vogogias, A., Preedy, K.F., and Milne, I. (2017). TetraploidSNPMap: Software for Linkage Analysis and QTL Mapping in Autotetraploid Populations Using SNP Dosage Data. Journal of Heredity 108, 438–442.

Haldane, J.B. (1930). Theoretical genetics of autopolyploids. Journal of Genetics 22, 359–372.

Koning-Boucoiran, C.F.S., Esselink, G.D., Vukosavljev, M., van’t Westende, W.P.C., Gitonga, V.W., Krens, F.A., Voorrips, R.E., van de Weg, W.E., Schulz, D., Debener, T., Maliepaard, C., Arens, P., and Smulders, M.J.M. (2015). Using RNA-Seq to assemble a rose transcriptome with more than 13,000 full-length expressed genes and to develop the WagRhSNP 68k Axiom SNP array for rose *(Rosa* L.). Frontiers in Plant Science 6, 249.

Li, X., Han, Y., Wei, Y., Acharya, A., Farmer, A.D., Ho, J., Monteros, M.J., and Brummer, E.C. (2014). Development of an alfalfa SNP array and its use to evaluate patterns of population structure and linkage disequilibrium. PLoS ONE 9, e84329.

McCallum, S., Graham, J., Jorgensen, L., Rowland, L.J., Bassil, N.V., Hancock, J.F., Wheeler, E.J., Vining, K., Poland, J.A., Olmstead, J.W., Buck, E., Wiedow, C., Jackson, E., Brown, A., and Hackett, C.A. (2016). Construction of a SNP and SSR linkage map in autotetraploid blueberry using genotyping by sequencing. Molecular Breeding 36, 41.

Preedy, K.F., and Hackett, C.A. (2016). A rapid marker ordering approach for high-density genetic linkage maps in experimental autotetraploid populations using multidimensional scaling. Theoretical and Applied Genetics 129, 2117–2132.

R Core Team. (2016). R: A language and environment for statistical computing. R Foundation for Statistical Computing, Vienna, Austria R version 3.3.2.

Revolution Analytics, and Weston, S. (2014a). doParallel: Foreach parallel adaptor for the parallel package. R package version 1.0.10.

Revolution Analytics, and Weston, S. (2014b). doSNOW: Foreach parallel adaptor for the snow package. R package version 1.0.14, 12.

Robins, J.G., Hansen, J.L., Viands, D.R., and Brummer, E.C. (2008). Genetic mapping of persistence in tetraploid alfalfa. Crop Science 48, 1780–1786.

Santos, J., Alfaro, D., Sanchez-Moran, E., Armstrong, S., Franklin, F., and Jones, G. (2003). Partial diploidization of meiosis in autotetraploid *Arabidopsis thaliana*. Genetics 165, 1533–1540.

Schmitz Carley, C.A., Coombs, J.J., Douches, D.S., Bethke, P.C., Palta, J.P., Novy, R.G., and Endelman, J.B. (2017). Automated tetraploid genotype calling by hierarchical clustering. Theoretical and Applied Genetics, 1–10.

Stam, P. (1993). Construction of integrated genetic linkage maps by means of a new computer package: Join Map. The Plant Journal 3, 739–744.

Stebbins, G.L. (1947). Types of polyploids: their classification and significance. Advances in Genetics 1, 403–429.

van Geest, G., Voorrips, R.E., Esselink, D., Post, A., Visser, R.G., and Arens, P. (2017a). Conclusive evidence for hexasomic inheritance in chrysanthemum based on analysis of a 183 k SNP array. BMC genomics 18, 585.

van Geest, G., Bourke, P.M., Voorrips, R.E., Marasek-Ciolakowska, A., Liao, Y., Post, A., van Meeteren, U., Visser, R.G., Maliepaard, C., and Arens, P. (2017b). An ultra-dense integrated linkage map for hexaploid chrysanthemum enables multi-allelic QTL analysis. Theoretical and Applied Genetics 130, 2527–2541.

Van Ooijen, J.W. (2006). JoinMap^®^ 4, Software for the calculation of genetic linkage maps in experimental populations. Kyazma B.V., Wageningen, The Netherlands.

Van Ooijen, J.W., and Jansen, J. (2013). Genetic mapping in experimental populations. (Cambridge University Press).

Voorrips, R.E., and Maliepaard, C.A. (2012). The simulation of meiosis in diploid and tetraploid organisms using various genetic models. BMC Bioinformatics 13, 248.

Voorrips, R.E., Gort, G., and Vosman, B. (2011). Genotype calling in tetraploid species from bi-allelic marker data using mixture models. BMC Bioinformatics 12:172.

Vos, P.G., Uitdewilligen, J.G.A.M.L., Voorrips, R.E., Visser, R.G.F., and van Eck, H.J. (2015). Development and analysis of a 20K SNP array for potato *(Solanum tuberosum):* an insight into the breeding history. Theoretical and Applied Genetics 128, 2387–2401.

Vukosavljev, M., Arens, P., Voorrips, R., Van’t Westende, W., Esselink, G.D., Bourke, P.M., Cox, P., Van de Weg, W.E., Visser, R.G.F., Maliepaard, C., and Smulders, M.J.M. (2016). High-density SNP-based genetic maps for the parents of an outcrossed and a selfed tetraploid garden rose cross, inferred from admixed progeny using the 68k rose SNP array. Horticulture Research 3, 16052.

Wu, R., Ma, C.-X., and Casella, G. (2002). A bivalent polyploid model for linkage analysis in outcrossing tetraploids. Theoretical Population Biology 62, 129–151.

Zheng, C., Voorrips, R.E., Jansen, J., Hackett, C.A., Ho, J., and Bink, M.C. (2016). Probabilistic Multilocus Haplotype Reconstruction in Outcrossing Tetraploids. Genetics 203, 119–131.

